# Glycoform Distributions and Heparan Sulfate Binding of Intact SARS-CoV-2 Spike Variants by DIA-ECCR Native Mass Spectrometry

**DOI:** 10.64898/2026.11.18.767578

**Authors:** Zhixin Xu, Chen Du, Eduardo Olmedillas, Erica Ollmann Saphire, Vicki H. Wysocki

## Abstract

Glycosylation influences the structure, immunogenicity, and evolutionary selection of the SARS-CoV-2 spike protein (S), but its extensive heterogeneity has hindered characterization of intact glycoform distributions. Here, we illustrate a data-independent analysis - electron capture charge reduction (DIA-ECCR) native mass spectrometry workflow for direct analysis of intact spike glycoforms across multiple variants. DIA-ECCR resolves variant-dependent differences in glycoform distributions and higher mass subpopulations. Monte Carlo simulations based on published site-specific bottom-up glycoproteomic data reproduce the central mass of the D614G experimental distribution and support a connection between intact and bottom-up measurements, while the broader experimental distributions and a second Gamma subpopulation indicate heterogeneity not captured by the simulations Mass photometry and charge detection mass spectrometry provide orthogonal measurements consistent with the DIA-ECCR results. DIA-ECCR also resolves mass shifts associated with binding of truncated heparan sulfate, demonstrating analysis of a heterogeneous protein:ligand complex. Together, these results outline a practical workflow that combines DIA-ECCR with native mass spectrometry (nMS) for characterizing intact glycoform distributions and ligand binding in highly heterogeneous glycoprotein assemblies, with potential applications in antigen characterization, vaccine design, and biopharmaceutical quality control.

## Introduction

Glycosylation, a common post-translational modification, plays a central role in protein folding and cell-cell signaling. Glycans on viral envelope proteins also contribute to viral entry and release.^1^ One example is the SARS-CoV-2 spike protein (S), a trimeric, heavily glycosylated surface protein that mediates viral entry into host cells.^2–4^ During the pandemic, several variants of concern (VOCs), including Alpha, Delta, Gamma, and Omicron, emerged with greater transmissibility and immune evasion than the prototypic Wuhan strain.^5–8^ Reduced vaccine efficacy against emerging SARS-CoV-2 variants is primarily driven by primary sequence mutations across the viral S, accompanied by site-specific differences in spike glycosylation that act as a structural shield against host immune recognition.^5,9^

The S is a trimer bearing approximately 22 N-glycosylation sites and up to 11 O-glycosylation sites per monomer.^6,10,11^ This extensive combinatorial glycosylation gives rise to a highly complex intact glycoform distribution, complicating intact-level characterization despite advances in site-specific glycan mapping. Accumulating evidence further shows that SARS-CoV-2 variants differ in site-resolved S glycosylation.^4,6,12–14^ At several N-glycosylation sites, including N165, N343, and N616, glycan compositions shift from high-mannose to complex forms.^15^ In contrast, O-glycans appear to change less across variants, although recombinant spike modifications may affect the T687 site.^6^ Together with sequence variation, these glycosylation changes can influence viral pathogenicity and entry mechanism. For example, Omicron’s altered entry pathway allows it to replicate primarily in the upper respiratory tract, underscoring the need for variant-specific structural characterization. ^16,17^

Native mass spectrometry (nMS) enables characterization of protein assemblies at the intact level, providing direct measurements of stoichiometry and topology while preserving noncovalent interactions.^18,19^ It is therefore well suited to large protein complexes and other heterogeneous biomolecular assemblies. However, in highly heterogeneous samples such as S and its VOCs, overlapping charge states often yield unresolved spectra, limiting further analysis.^11,20^ Glycosylation contributes substantially to this heterogeneity through variable site occupancy and glycan composition, further complicating intact-level analysis.

Single particle methods and charge manipulation strategies have emerged as effective approaches for characterizing highly heterogeneous glycoproteins.^21–29^ Single particle methods, including charge detection mass spectrometry (CDMS) and mass photometry (MP), have been used to measure the intact mass of S produced in different cell lines, estimate surface glycosylation coverage after PNGase F treatment, and assess the binding avidity of intact S to monoclonal antibodies.^20,30,31^ In parallel, charge manipulation approaches offer an alternative route to simplifying complex spectra. For example, proton transfer reactions (PTR) have been used to probe S interactions with ACE2 and heparin binding.^26^ Data-independent acquisition-proton transfer charge reduction (DIA-PTCR) has likewise shown that heterogeneous glycoform distributions can be resolved for more specific profiling of glycosylated biotherapeutics.^27^ More recently, Shaw et al. and subsequently Le Huray et al. illustrated electron capture charge reduction (ECCR), and Shaw et al. showed that ECCR can resolve overlapping charge states of S, enabling deconvolution and measurement of its intact mass distribution.^28,29^ Nevertheless, workflows that map intact glycoform distributions across multiple spike variants and directly connect those measurements to site-specific glycoproteomics remain limited.

In this work, we used DIA-ECCR to characterize intact glycoform distributions across multiple S variants. The resulting distributions were compared with Monte Carlo simulations based on glycoproteomic studies in literature. More broadly, this study advances the use of nMS for measuring intact glycoform distributions of highly heterogeneous glycoproteins and supports DIA-ECCR as a complementary approach to bottom-up glycoproteomics. DIA-ECCR also resolves mass shifts associated with the binding of truncated heparan sulfate. Together, these results establish DIA-ECCR as an effective strategy for characterizing intact glycoform distributions in highly heterogeneous systems. The workflow should be useful for rapid characterization of glycosylated antigens and for quality control in biopharmaceutical production.

## Results

### DIA-ECCR reveals features of the intact glycoform distribution

The VFLIP D614G spike (S), a construct stabilized by five (V) proline substitutions, a flexible cleavage-site linker, and an inter-protomer disulfide bond, is extensively glycosylated, introducing substantial heterogeneity, yielding a broad, unresolved *m/z* signal at 10-14k (Figures 1A inset/1D).^11^ By isolating a narrow *m/z* window (100 *m/z*) and then performing ECCR, overlapping signals within that window were simplified by reducing the charge, thereby spreading the peaks more effectively in *m/z* space and facilitating deconvolution (Figure 1B). This method has been shown to simplify overlapping signals and improve characterization of glycoform distributions for S, thyroglobulin, and antibodies.^29,32^ As shown in Figure 1C, deconvolution of the 11,900– 12,000 *m/z* isolation window yielded six distinct peaks in the zero-charge mass spectrum, grouped into prominent mass intervals averaging 12,030 Da. Because mass-to-charge ratio (*m/z*) scales mass with charge state (mass ≈ z × [*m/z*]), isolating a narrow window around ∼12,000 m/z means that ions differing by one charge state (z vs. z+1) must differ in neutral mass by approximately 12,000 Da. The observed ∼12 kDa mass gaps thus directly correspond to species that co-isolated in the same *m/z* window but possessed adjacent charge states in the unresolved spectrum, demonstrating that charge state overlap—in addition to structural heterogeneity—drives the broadness of the original signal.

**Figure 1.**
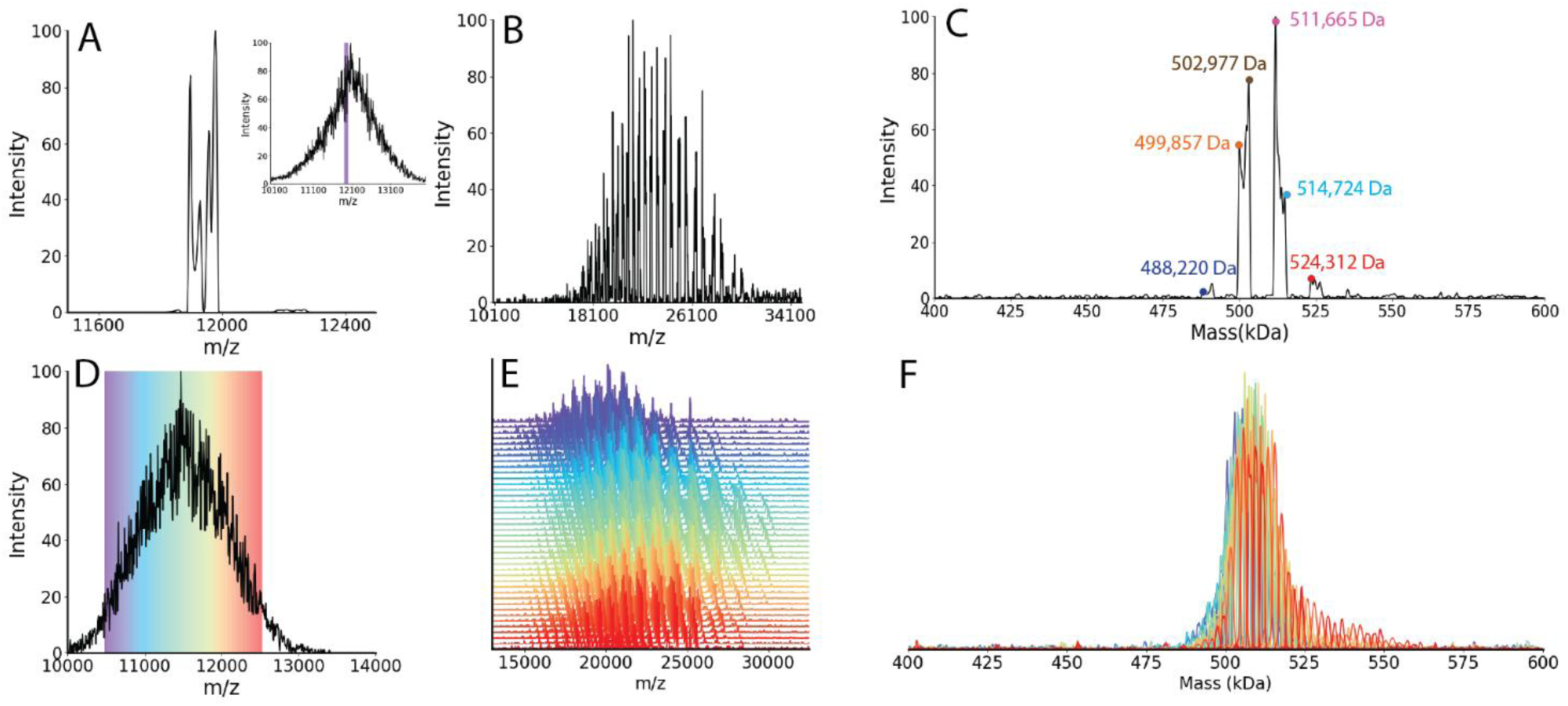
Workflow for DIA-ECCR to resolve intact glycoform distributions of the S. (A) Isolated 100 m/z window from D614G S (11900-12000 m/z). The inset shows the full distribution and the selected window. (B) ECCR at 7 V applied to the isolated 100 m/z window. (C) Deconvolved spectrum from panel B. (D) Zoomed-in spectrum for the D614G S. Colors are labeled according to panel E, with each color representing a 50 m/z isolation window. (E) ECCR at 7 V applied to each isolation window from panel D. (F) Overlaid deconvolved spectra from panel E. Colors are the same as in panels D and F.

Building upon the ECCR isolation method previously described,^29^ we establish a workflow to reconstruct the mass distribution of complex glycoforms. By selecting sequential 50 *m/z* window selection across a portion of the main peak (11,075-11,425 *m/z*) reconstructed a mass distribution of 487-526 kDa using a 5% intensity cutoff. When repeating the 50 *m/z* isolation window across five consecutive steps, we observed a cyclic mass shifting pattern in which the dominant species progressively shifts and then returns to the lowest mass species. To further confirm this distribution, 41 spectra were collected using 50 *m/z* windows with ECCR across 10,000-12,000 *m/z* (Figure 1E). Lower signal intensity at the edges of the *m/z* range may lead to missing lower charge state peaks. Overlaying these spectra produced a cumulative mass distribution (Figure 1F). Although there is a slight shift toward a higher mass from the purple to the red tracings, the overall mass envelope remains relatively consistent. As a control, partial deglycosylation with PNGase F altered the glycoform distribution (Figure S1). After one day of incubation with PNGase F, a large amount of glycosylation remained on the S, which is also observed in Stiving et al. experiments.^30^

### Comparison of different VOCs using the DIA-ECCR workflow

After establishing and validating the DIA-ECCR workflow for a representative variant, D614G, we next applied it to multiple variants to demonstrate its ability to resolve mass distribution differences. A quick examination of eight variants, all expressed in CHO cells in the same laboratory, shows that the mass distribution for single-slice spectra varies across variants (**Error! Reference source not found.**). The mass range for all the variants is around 500 kDa, with different peak widths and different global mass distributions. Specifically, BA.4/5 has the narrowest mass distribution, and the Gamma variant has the broadest mass distribution among these eight variants, suggesting Gamma is a more heterogeneous sample than the others. Although BA.1, BA.2, and BA.4/5 belong to the same sublineage - Omicron, the potential mass distribution for BA.1 is wider than the others. To fully understand the actual differences between these variants and to examine the limits of the DIA-ECCR method, a more comprehensive characterization of four selected VFLIP variants was conducted for Omicron BA.1, Omicron BA.4/5, D614G, and Gamma (Figure 2B).^11^ D614G is an early variant with a one-amino-acid substitution, while Omicron and Gamma are two VOCs recognized for their high infection rates during the pandemic.^8,10^ The trimer protein sequence masses (without glycosylation) are 406 kDa for the D614G variant, 406 kDa for the Omicron BA.1 variant, 407 kDa for the Omicron BA.4/5 variant, and 407 kDa for the Gamma variant.

**Figure 2:**
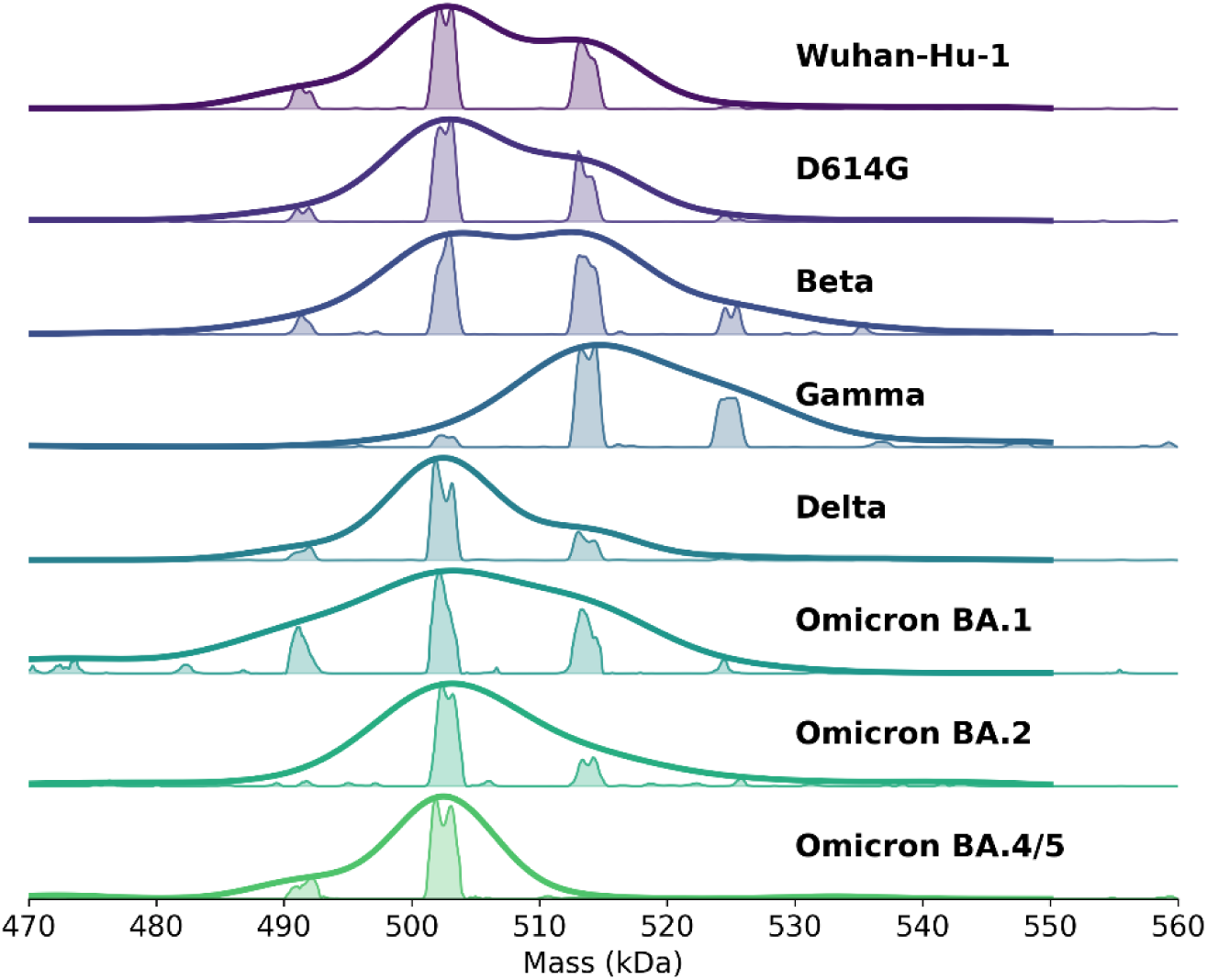
Deconvolved mass distributions from single-slice measurements for the eight variants indicated. Kernel density estimation was applied to estimate the potential mass distribution of those variants.

The sequence mutations do not significantly change the intact mass of the Gamma variant compared with other variants. While many glycoproteomics studies have shown that glycosylation of different S variants is altered, the effect at the intact level has not yet been evaluated.

The conventional ensemble mass spectra (acquired without dissociation or ECCR) were obtained for those four variants. After applying ECCR 7 V, BA.4/5 showed the most discrete charge state peaks (Figure S2). The deconvolved masses after applying ECCR 7 V indicate that the glycosylated Gamma variant has a higher intact mass than other variants. Previous glycoproteomics studies have shown that Gamma has two additional N-glycosylation sites compared to other variants, which partially explains the increased glycoform masses.^6,15^ Similarly, Robinson and coworkers showed that artificially constraining glycan heterogeneity to a uniform high-mannose type enabled discrete charge state resolution of intact S in mass spectra.^33^ Their results and our ECCR results suggest that some of the spike’s spectral heterogeneity can be attributed to glycan microcomplexity. As a result, the simpler mass spectrum of D614G and BA.4/5 reflects a lower degree of glycan microheterogeneity across the intact S trimer.

Reconstructed DIA-ECCR mass distributions reveal distinct variant-dependent profiles across the four spike constructs analyzed in greater detail (Figure 3). Although D614G and BA.1 have similar sequence mass and mass distribution centers, the mass distribution peak width for D614G is narrower than that of BA.1. Reconstructed DIA-ECCR mass profiles for the Gamma variant reveal a primary distribution centered at 518 kDa (FWHM = 16.7 kDa) alongside a distinct, low-abundance higher-mass envelope spanning 540-600 kDa (centered at 554 kDa, FWHM = 50.8 kDa; Figure 3B). The dominant mass species in the primary distribution for Gamma is higher than in the D614G and BA.4/5 variants, matching the single-slice measurements. Crucially, the ∼36 kDa mass shift defining the secondary subpopulation is far too small to represent an additional protomer or higher-order oligomeric state, yet significantly exceeds routine salt adduction, so it is likely glycan-related.

**Figure 3:**
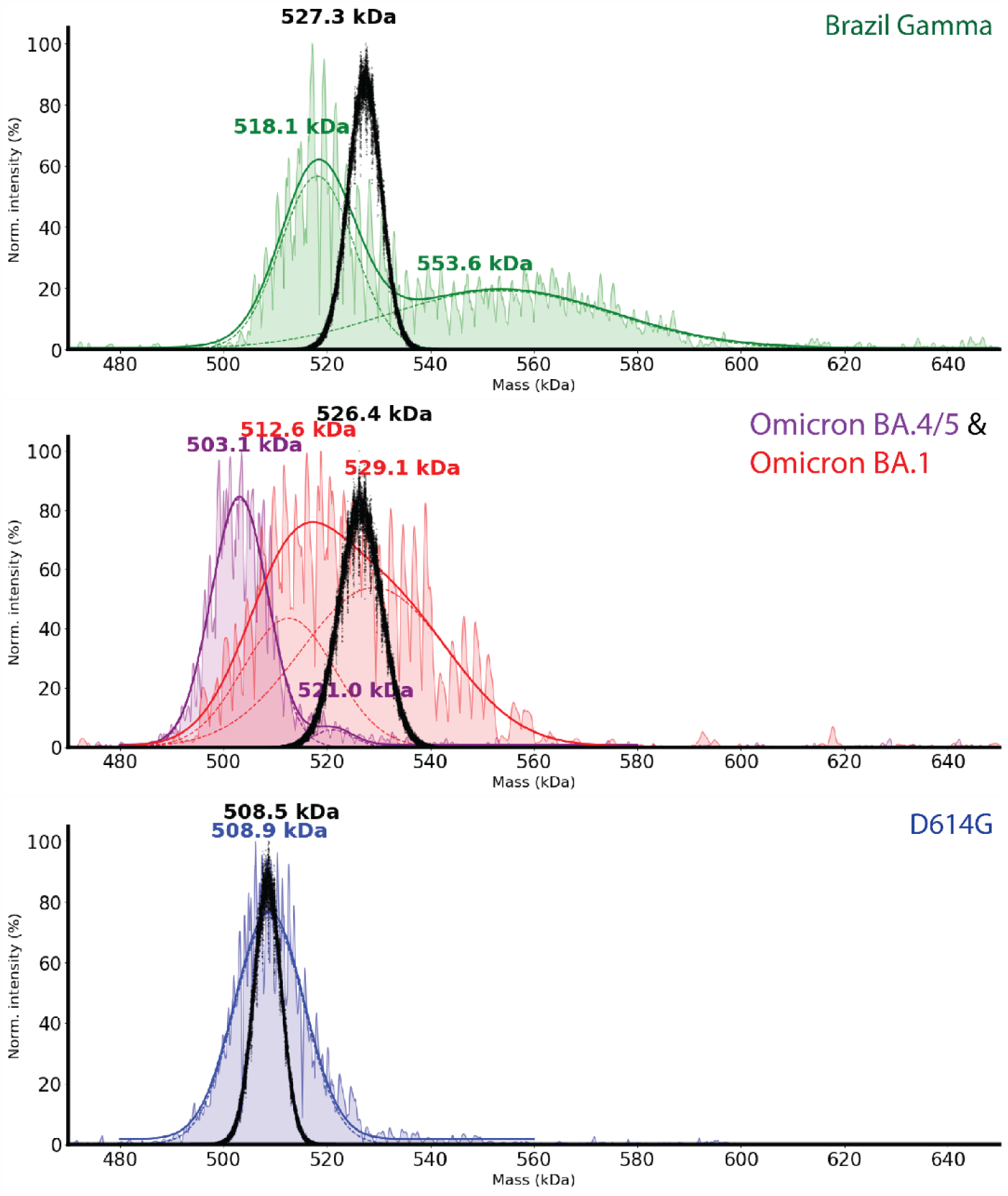
Full mass distribution envelopes for four selected variants from the ECCR method (colored lines). The displayed profile represents the combined data from 10 slices, using the maximum intensity across slices at each x-position. Three simulation results from glycoproteomics data (black dots). Individual Gaussian fits for different VOC are shown as dashed lines, and the solid line represents the overall fit obtained by summing the Gaussian components. The center mass for individual gaussian fits were reported.

To orthogonally validate DIA-ECCR ensemble results, we benchmarked our measurements against established MP and CDMS techniques, (Figure 4). The three experimental methods agree reasonably well across all tested variants, consistent with previous cross-technique comparisons for large heterogeneous samples.^29,34,35^ Specifically, MP and CDMS confirmed the lower mass averages and narrower mass distributions of D614G and Omicron BA.4/5 relative to other variants. For heterogeneous variants, ECCR and CDMS resolved a broad mass distribution in Omicron BA.1, whereas all three techniques consistently detected a low abundance, high mass subpopulation in Gamma. The overall alignment across these independent methods confirms that the observed mass variations represent biological heterogeneity rather than method specific artifacts.

**Figure 4.**
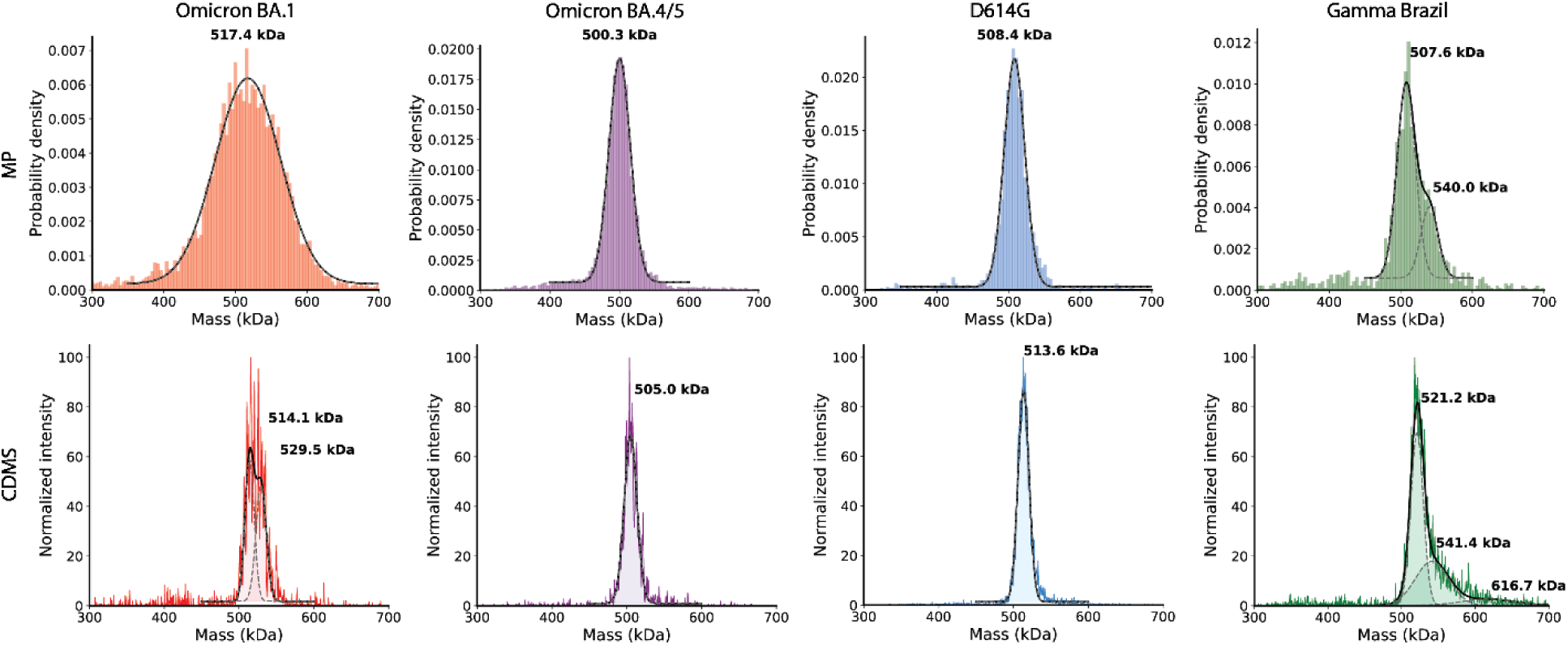
Single particle methods for S VOC. The top panel was collected by MP, and the bottom panel by CDMS. Different colors refer to different VOCs. Gaussian fits were applied to the mass distributions for MP and CDMS results. Individual Gaussian fits are shown as dashed lines, and the solid line represents the overall fit obtained by summing the Gaussian components.

### Modeling Spike Glycoform Distributions via Monte Carlo Simulations

To contextualize the experimental nMS results relative to bottom-up estimates, we modeled hypothetical intact S mass distributions for three VOC (Gamma, Omicron BA.1, and D614G) via Monte Carlo (MC) simulations using published site-specific glycoproteomics data, assuming statistically independent site occupancy.^4,10,12,36^ To ensure sampling sufficiency, an extensive number (10×10^7^) of MC simulations was performed, and the convergence of the center mass was observed. (Figure S3) The simulated mean mass (508 kDa) of D614G aligned closely with the experimental DIA-ECCR value (509 kDa) (Figure 3). However, the experimental data mass range (487-526 kDa, above 5 % signal intensity) was broader than the simulated range (493-522 kDa, above 5 % relative frequency Figure 2B). The broader D614G experimental distribution may reflect inter-site glycan dependencies, including cooperative or sterically constrained processing at adjacent glycosylation sites. Because proteolysis in bottom-up workflows removes information about which glycans coexist on the same intact molecule, DIA-ECCR may provide complementary evidence of glycan crosstalk that is not accessible from peptide-level measurements alone. For the Gamma variant, although the Monte Carlo simulation (centered at 527 kDa, FWHM = 7.58 kDa) overlaps both the lower and higher mass envelopes of Gamma, the simple model can’t predict the two mass distributions seen in the nMS. For the Gamma higher mass subpopulation obtained from ECCR experiments, the mass increment points to two primary possibilities. First, it may reflect trimers incorporating hyper-processed monomers with extended complex branching or elevated sialylation, consistent with previous reports of unequal monomer processing within intact spike assemblies.^20^ Second, it could indicate distinct trimer subpopulations governed by correlated, non-random glycan occupancy, reflecting a level of meta-heterogeneity that inherently escapes site-averaged bottom-up measurements and independent Monte Carlo models. A clear mass shift between the intact glycoform profiles of BA.1 and BA.4/5 indicates considerable glycosylation differences even within the Omicron sublineage. Because site-specific glycosylation data for BA.4/5 are currently unavailable, Monte Carlo glycoform simulations were not performed for direct comparison. This limitation highlights the need for variant-specific, site-specific glycoproteomic mapping to accurately model intact mass distributions.

Beyond potential site-to-site crosstalk, several nonexclusive technical and methodological factors may also contribute to the observed mass dispersion. (1) Although prior studies suggest a total of 22 N-glycan sites and up to 11 O-glycan sites, the bottom-up mass spectrometry data in the publication identified only 20 N-glycan sites and 8 O-glycan sites for the D614G variant, underreporting glycosylation microheterogeneity for intact S.^6,11,12,36^ Because O-glycosylation occupancy is low, the effect of these missing O-glycans is likely small. (2) The glycoproteomics data were obtained from proteins expressed in Human Embryonic Kidney (HEK) 293 cells, whereas the proteins analyzed here by experiments were expressed in Chinese Hamster Ovary (CHO) cells. Robinson and Jarrold reported similar results: CHO cell-expressed proteins tend to have a narrower distribution than proteins expressed in HEK.^20^ (3) Bottom-up proteomics remains subject to analytical biases, including differential peptide ionization efficiencies and dynamic range limitations that obscure low-abundance glycoforms.

### Characterization of S binding to heparan sulfate

Heparan sulfate (HS) is a linear polysaccharide of repeating disaccharide units (glucosamine and hexuronic acid) that is normally found on most animal cell surfaces or in the extracellular matrix.^37–41^ Many viruses, such as the influenza virus, herpes simplex virus (HSV), and human immunodeficiency virus (HIV), use glycoconjugates on the host cell surface as attachment sites, facilitating infection.^41,42^ Clausen et al. show that HS influences SARS-CoV-2 infection across different cell types, and that HS binds to S and ACE2 to form a tertiary complex.^38^ This established interaction provides a relevant heterogeneous protein:ligand system for testing whether DIA-ECCR can resolve ligand-associated mass shifts.

Spike binding to HS is difficult to study experimentally because HS is a mixture of various disaccharide chains, each containing around 150 units. Herein, due to the heterogeneity of HS and spike, a truncated version of HS, dp20 (average mass from 8k-10k Da), has been used to test ECCR’s ability to monitor binding. After binding to HS, the mass spectra show no significant differences (Figure 5), possibly because S and S bound to HS have very similar structures and thus similar charge state distributions. By selecting the entire unresolved envelope for the S:HS complex and applying ECCR, the charge reduced spectrum shows evidence of binding, where resolved charge states can be obtained for spike, while the charge state distribution stays unresolvable for S-HS complexes. A single slicing window revealed that the spike’s detailed mass distribution has widened. Surprisingly, the unbound S still appears in deconvolved mass spectra. Despite the small mass difference (∼10.3 kDa), the S-HS complex shows lower charge reduction efficiency, even though the original charge distributions are similar (bound complex: 46-41; unbound complex: 47-41). (Figure S4) This suggests that the HS reacts with the positive groove of the spike but does not introduce significant restructuring in the spike trimer. After applying DIA-ECCR to the S-HS complex, the Gaussian fit shows four new mass species in the full mass distribution, suggesting that at least four copies of HS bind to the spike. These results suggest that ECCR can be used to characterize small mass shifts in ligand binding across heterogeneous complexes. Given that the centroid positions of the four mass species differ by similar amounts, the results strongly imply that HS has low binding selectivity, as previous studies have shown.^43,44^ Meanwhile, because the fourth mass distribution is less abundant, S-HS binding is difficult to saturate even at a 1:10 concentration ratio. When high-voltage HCD was applied, the bound HS was ejected, resulting in a narrow mass distribution similar to that of an unbound S (Figure S5). This gas-phase ejection confirms that HS interacts with S via non-covalent mechanisms, mirroring previous nMS observations by the Kaltashov group using the related glycosaminoglycan heparin and RBD of spike.^26^ All in all, this study has shown that ECCR can be used to characterize a highly glycosylated immunogenic target, such as S, that binds a small ligand, thereby validating its glycoprotein-ligand interactions.

**Figure 5:**
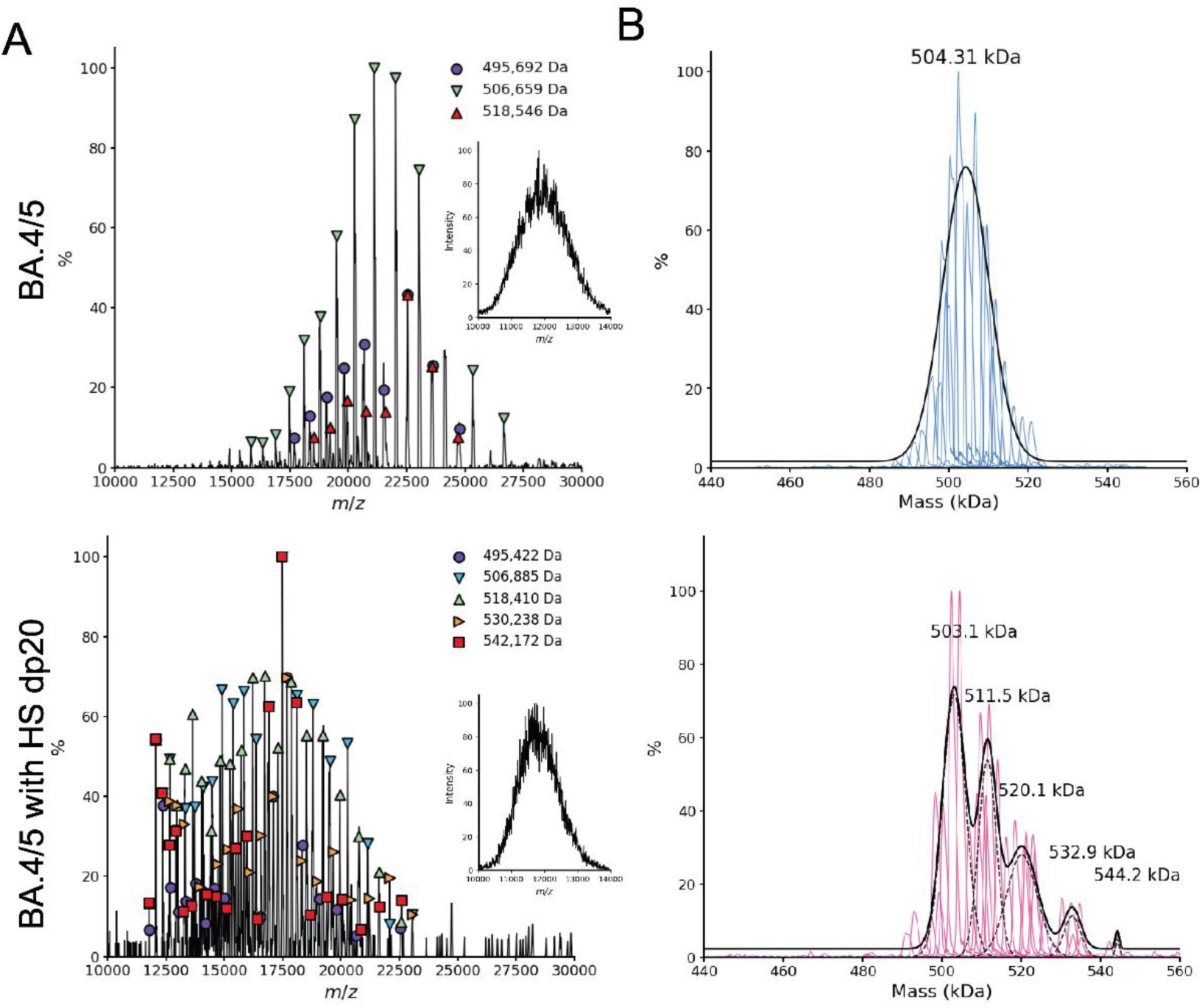
Comparison of BA.4/5 without (top) and with (bottom) bound heparan sulfate dp20. (A) ECCR applied to the 11775-11825 m/z isolation window. Insets are the fly-through spectra for BA.4/5 and BA.4/5 with HS dp20. (B) Reconstructed mass distribution for BA.4/5 and HS dp20.

## Discussions and Conclusion

To tackle the complexity of heterogeneous glycoproteins, we developed a DIA-ECCR profiling approach. A rapid screening of eight initial variants let us map the glycoform landscape and choose four variants for closer study. We then compared the DIA-ECCR results from three variants with MC simulations based on bottom-up experiments, and all three variants showed wider glycoform distributions than the simulations predicted. These discrepancies between bottom-up predictions and intact nMS mass are beginning to be recognized as new nMS techniques have been developed.^20,32,34^ They arise mainly because bottom-up workflows struggle to give unbiased, quantitative glycan occupancies. Researchers have begun to address this by feeding experimental intact nMS intensity distributions directly into stochastic simulations, a promising route to more accurate models of intact glycoform landscapes.^34^

Previous molecular dynamics studies have shown that glycosylation changes are accompanied by structural alterations: Omicron S shows altered glycan shielding and greater solvent accessibility, along with increased ACE2 binding, relative to wild-type.^49^ However, site-specific comparisons within the same lineage have not yet been reported.^15,45^ Instead, most studies have focused on comparisons between variants and their ability to interact with mAbs and ACE2 receptors.^6,15,46–48^ These observations evaluated glycosylation composition differences between variants but not differences within Omicron sublineages. Although later Omicron variants like BA.2.12.1, BA.4, and BA.5 belong to the same Omicron sublineage, they differ enough to show escape from antibody neutralization compared to previous Omicron subvariants,^50^ highlighting the need for variant-resolved characterization of S. In this work, we compared the two Omicron sublineage variants BA.1 and BA.4/5, which showed distinct intact glycosylation distributions. This suggests that intra-lineage sequence mutations and local structural changes are sufficient to modulate macro-level glycan heterogeneity, and high-resolution, site-specific comparisons across sublineages are an important future direction for determining how local microenvironments dictate regional glycan processing.

By integrating DIA-ECCR with native mass spectrometry, this study establishes a robust framework for disentangling complex glycoform heterogeneity in intact heterogeneous glycoproteins. Initial rapid profiling across eight Spike variants mapped the baseline landscape. This work then demonstrates DIA-ECCR’s capability of resolving intact glycoform distributions of four VOCs of the S complex. The center mass of the dominant Gaussian fits for these four VOCs (D614G, BA.1, BA.4/5, and Gamma) are 509 kDa, 529 kDa,503 kDa, and 518 kDa, respectively. The results align well with other parallel techniques (MP, CDMS) and the Monte Carlo simulations based on bottom-up proteomics. While DIA-ECCR successfully captures system-level glycan heterogeneity, one possible explanation for the differences between the simulation and experimental datasets is complex glycan coexistence. ECCR was also used to probe the interaction between two heterogeneous samples: spike and heparan sulfate (HS). The results show that after spike binds to HS, four new mass envelopes, with spacing consistent with the mass of HS, are observed in the intact glycoform distributions. Overall, combining DIA-ECCR with native MS provides a powerful framework for mapping the overarching glycoform macro-heterogeneity of complex glycoprotein assemblies. Looking forward, pairing these intact mass spectra with advanced computational deconvolution and stochastic modeling algorithms will enable the extraction of higher-order inter-site glycan correlations, revealing co-processing dependencies that remain invisible to site-averaged bottom-up proteomics.

## Methods

### Material and Sample Preparation

VFLIP S and VOCs were expressed and purified as previously described by the Ollmann Saphire lab.^11^ The sequences of D614G, BA1, and Gamma were obtained from the Saphire lab. C-reactive protein (CRP) and GroEL were used as calibrants for molecular mass calibration in CDMS, whereas beta-amylase (BAM) and thyroglobulin (TG) were used for mass photometry; all were purchased from Sigma-Aldrich (St. Louis, MO). Heparan sulfate dp20 was purchased from Galen Molecular (North Haven, CT). PNGase F was purchased from Promega (Fitchburg, WI). For glycan removal, S were mixed with PNGase F in a 1:4 (µg of protein: umol of enzyme) ratio and incubated at 37 ℃ for 1 days. VFLIP S and VOCs were buffer exchanged into 200 mM ammonium acetate (Sigma-Aldrich) using a 50 kDa or 100 kDa molecular weight cutoff (MWCO) Amicon Ultra centrifugal filter (Sigma-Aldrich). CRP and GroEL were buffer-exchanged into 200 mM ammonium acetate using Bio-Rad Micro BioSpin P-6 columns. VFLIP VOC samples for nMS and nMS-CDMS were analyzed at complex concentrations of 500 nM to 1 µM of complex, whereas samples for mass photometry were analyzed at 10-25 nM.

### Mass Spectrometry

All mass spectrometry experiments were performed on a Thermo Q Exactive Ultra-High Mass Range Orbitrap mass spectrometer (Thermo Fisher Scientific) equipped with a custom Electron-based Dissociation-Surface Induced Dissociation (ExD-SID) device developed in a research collaboration with the former eMSion, Inc. (Corvallis, OR).^29^ The ExD-SID device was installed between the quadrupole mass filter and the C-trap in place of the transfer multipole. The UHMR quadrupole was used for narrow window mass-to-charge (*m/z*) selection, and ECCR was used to resolve overlapping signals within the selected *m/z* range. Narrow window selection refers to quadrupole isolation of a limited *m/z* range (in this case, 30-100 *m/z*), narrower than the full heterogeneous distribution, which spans approximately 4000 *m/z*. Electron capture uses electrons generated by a heated filament to reduce charge states. ECCR voltages were described previously.^29^ A Sutter P-97 micropipette puller was used to prepare tips for nanoelectrospray ionization from uncoated borosilicate capillaries (O.D. 1 mm, I.D. 0.78 mm) using in-house optimized programs. The spray voltage typically ranged from 900 to 1100 V. For all experiments, the inlet temperature was 250 ℃, in-source trapping was set to −50 V, and the maximum injection time was 50-3000 ms. CDMS experiments were performed at a resolution of 200 k at *m/z* 400, with IST set to −50 V and trapping gas 2 enabled in Direct Mass Technology mode (Thermo Fisher Scientific). The CDMS experiment was calibrated using CRP (5-mer) and GroEL (14-mer and 13-mer ejected at high IST).

ECCR data were analyzed using the deconvolution software UniDec.^51^ For spectra acquired without ECCR, the following deconvolution parameters were used: charge range, 1-60; mass range, 400-600 kDa; sample mass every (Da), 1; peak FWHM (Th), 30-200; beta, 50.0; charge smooth width, 1.0; point smooth width, 10.0; mass smooth width, 0.0; maximum number of iterations, 100; peak detection, 5000 Da; and peak detection threshold, 0.5. For data acquired with narrow quadrupole *m/z* selection and ECCR, the deconvolution parameters were: charge range, 1-60; mass range, 400-600 kDa; sample mass every (Da), 1; peak FWHM (Th), 5.0; beta, 0; charge smooth width, 1.0; point smooth width, 1; mass smooth width, 0; maximum number of iterations, 100; peak detection, 500 Da; and peak detection threshold, 0.1. Deconvolved spectra were compared with glycoproteomic data using Monte Carlo simulations.^12,36^ CDMS data (Direct Mass Technology^TM^) were first analyzed using STORIboard software (Thermo Fisher Scientific) and then with UniDecCD.^21,52^ The UniDecCD processing parameters were: charge range, 35-55; *m/z* bin size, 2-6; mass bin size, 2-6; *m/z* spread FWHM, 2-6; charge spread FWHM, 2-6; and point smooth, 1.

The Monte Carlo simulation was carried out using in-house Python code. Site-specific glycan compositions and occupancy percentages were obtained from published glycoproteomics studies.^4,6,12,36^ The D614G simulation included 28 glycosylation sites per monomer: 20 N-glycan sites and 8 O-glycan sites. ^12,36^ The Gamma simulation included one additional N-glycan site, and the Omicron BA.1 simulation included two additional N-glycan sites.^4,6^ At each site, a reported glycan composition was sampled according to its occupancy-weighted probability to calculate the glycan mass of a spike monomer. The sequence masses of three monomers and three independently sampled glycan masses were then summed to obtain each simulated trimer mass. The equation used to calculate one possible total trimer mass is shown below:

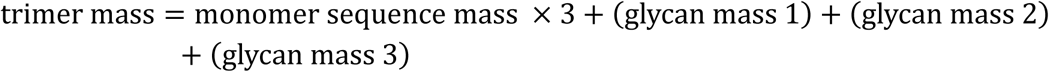

Glycan masses 1, 2, and 3 each represent the sum of randomly selected glycans across all 28 glycosylation sites in each monomer. These three glycan masses may differ because the selected glycan compositions at the 28 sites are not necessarily the same for each monomer within the spike trimer. To verify that the chosen number of Monte Carlo iterations is sufficient, we performed the simulation with increasing sample sizes, from 5 × 10^4^ to 1 × 10^7^ individual runs (Figure S2). As the number of iterations increased, the estimated mean mass converged to a stable value, with negligible differences between the larger sample sizes. This convergence indicates that the iteration count used (5×10^6^) in this report is sufficient to obtain a reliable estimate and that further increasing the number of runs would not meaningfully change the result. 5 × 10^6^ times were used to report the results. For faster analysis, the final trimer masses were rounded to integers, and the resulting distribution was analyzed by counting the occurrence of each glycoform.

### Mass Photometry

A Refeyn TwoMP instrument was used for all mass photometry experiments. The CultureWell reusable gasket and high-precision microscope glass slides (24 × 50 mm) were cleaned and placed on the instrument for mass measurements. BAM (56 kDa monomer, 112 kDa dimer, and 224 kDa tetramer) and thyroglobulin (335 kDa monomer and 670 kDa dimer) were used as calibrants (Sigma-Aldrich).^29^ All data were collected for 1 min using AcquireMP software and then analyzed with DiscoverMP software. Gaussian fitting was applied to MP histograms to determine masses. All MP measurements were performed by droplet dilution in 200 mM ammonium acetate solution (Sigma-Aldrich).

## Acknowledgement

The authors thank Regina M. Edgington and Steffen Lindert, and his group members for helpful suggestions on the Monte Carlo simulations. The authors gratefully acknowledge funding from the National Institutes of Health awarded to Jared B. Shaw. (R43GM140749 (SBIR for Shaw/Wysocki group development of ExD-SID device), E.O.S. (P01AI165072), and V.H.W. (P41GM128577 and RM1GM149374).

## Author Contributions

**Z.X.:** Investigation, Methodology, Conceptualization, Visualization, Software, Formal analysis, Writing – original draft, review and editing. **C.D.:** Investigation, Methodology, Conceptualization, Writing – review and editing. **E.O.:** Resources, Writing – review and editing. **E.S**.: Funding Acquisition, Writing – review and editing. **V.W.:** Funding Acquisition, Conceptualization, Writing – review and editing.

