## supplemental information for "Glycoform Distributions and Heparan Sulfate Binding of Intact SARS-CoV-2 Spike Variants by DIA-ECCR Native Mass Spectrometry"

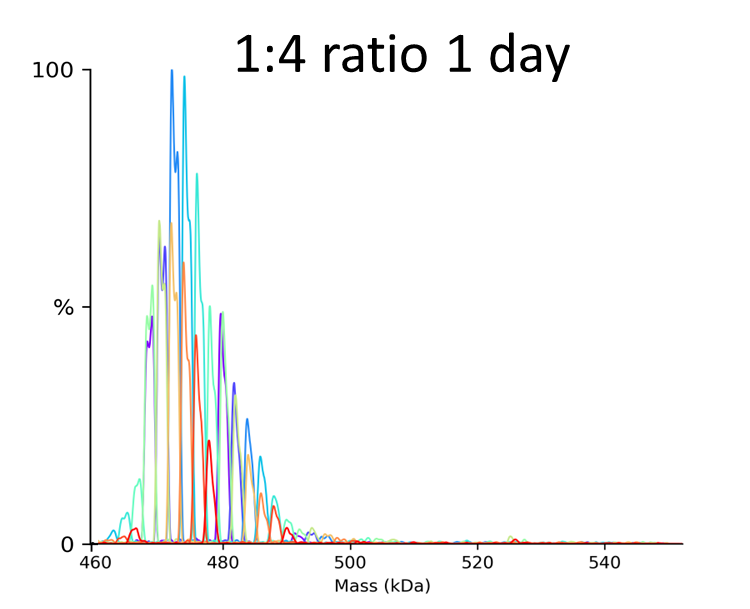

Figure S1: PGase F deglycan on the D614G spike protein. The glycoform distribution was obtained using the DIA-ECCR workflow described in the main text.

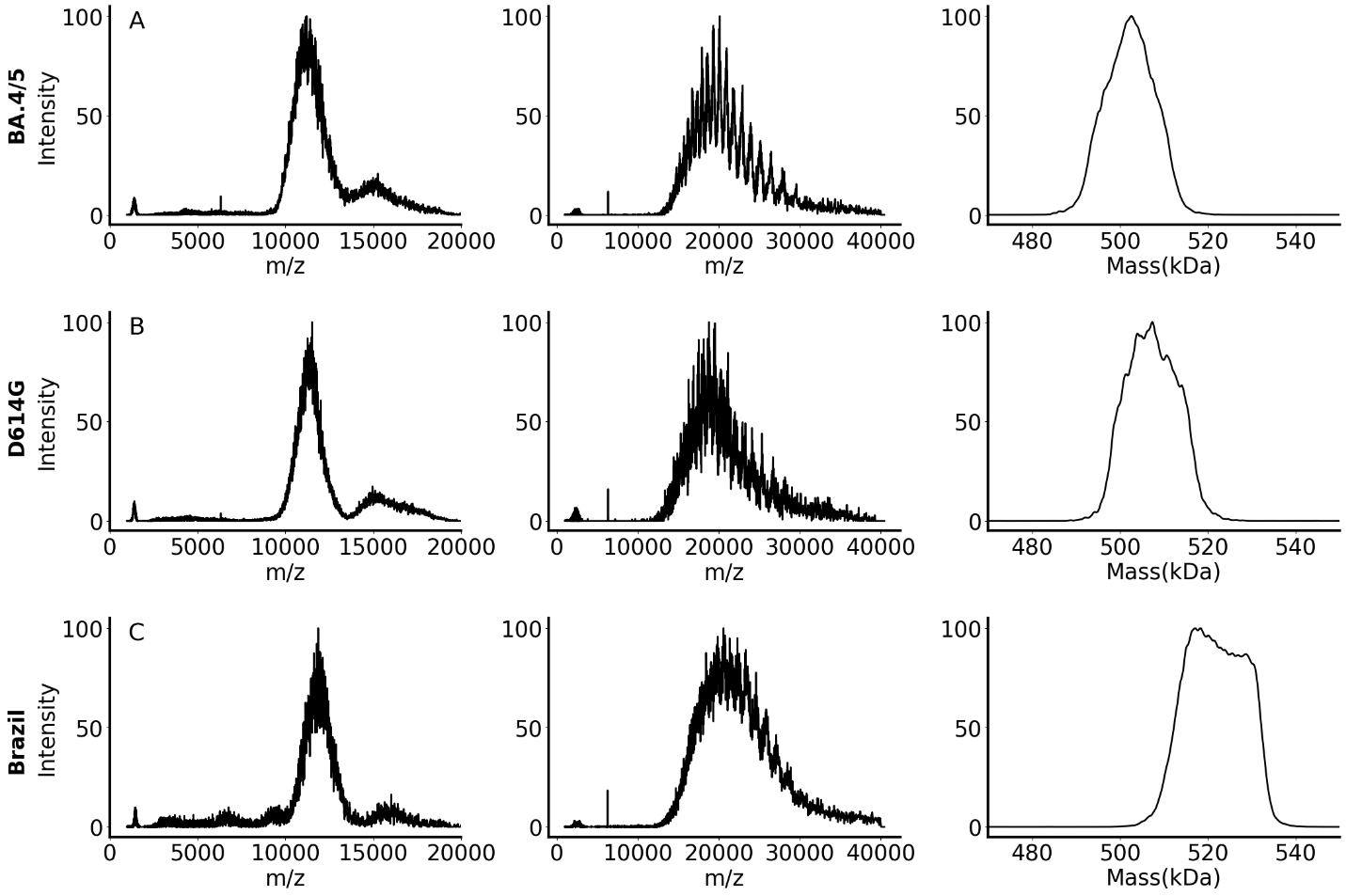

Figure S2: Comparison between four VOC. The first column refers to the flythrough spectrum for three different variants, and the second column displays the full distribution of spikes(isolation window 10-14k m/z) with ECCR; the third column shows the deconvolved mass spectra from the second column. (A) BA.4/5 variant: 502,641 Da. (B) D614G variant: 507,322 Da. (C) Brazil variant: 514,739 Da.

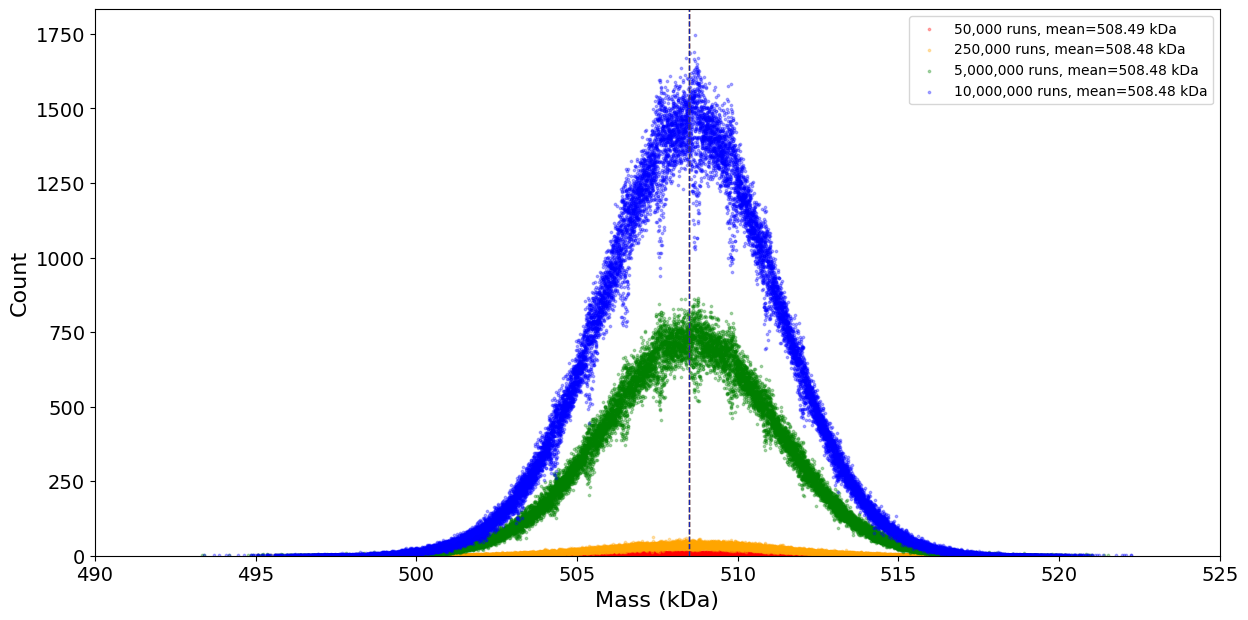

Figure S3: Different counts of Monte Carlo simulations for D614G spike protein. The center mass for this simulation has converged to 508.5 kDa.

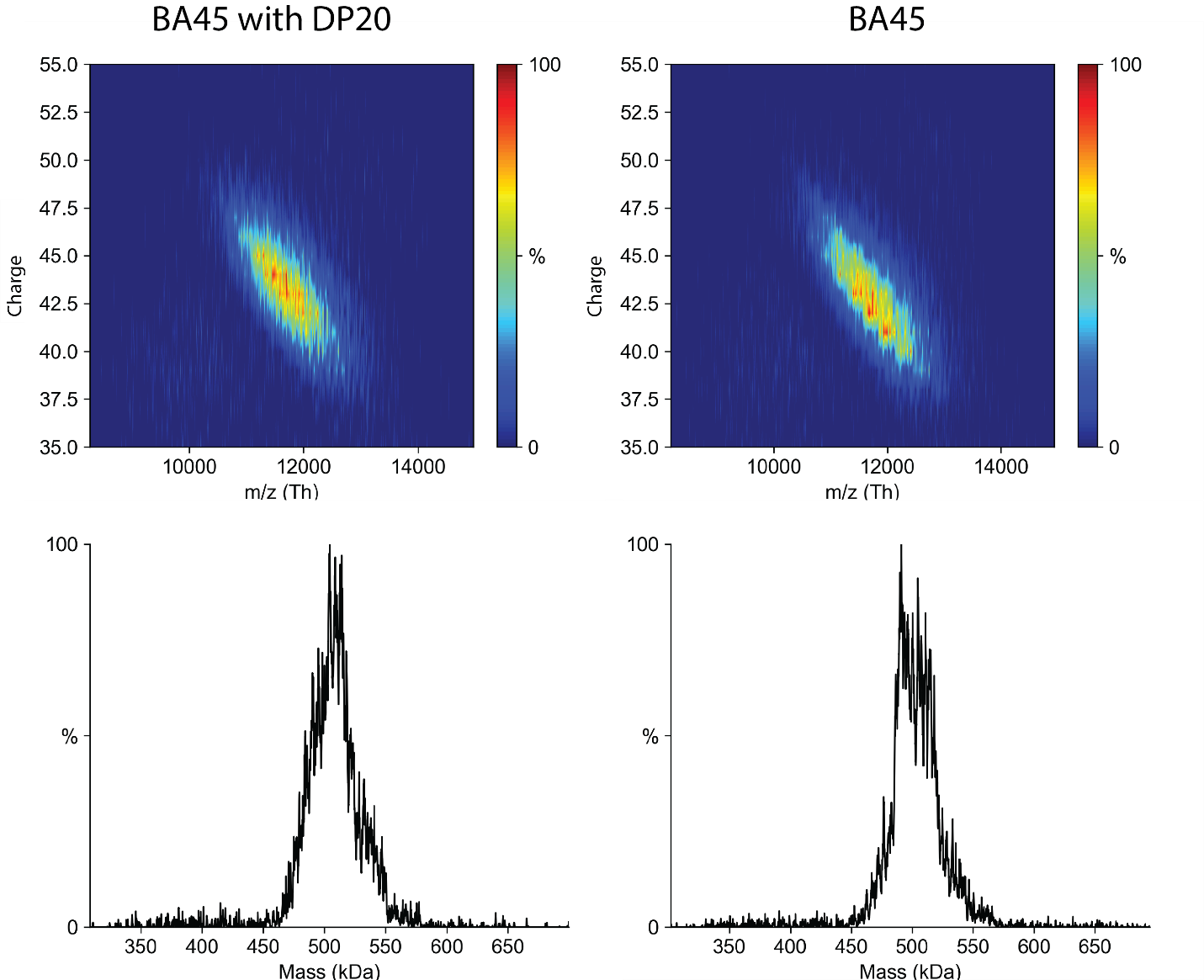

Figure S4: CDMS for BA45 variant bind and without bind DP20.

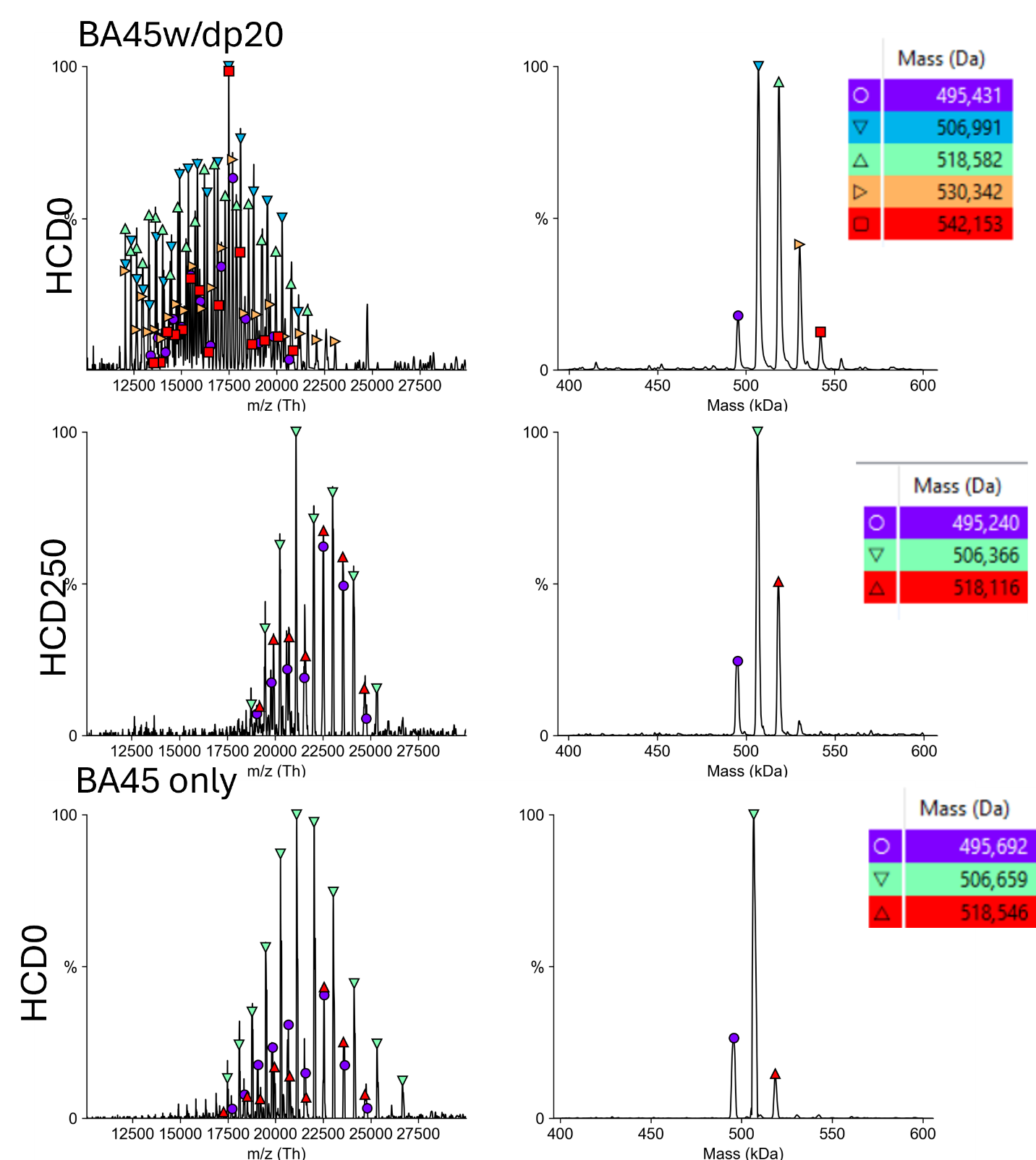

Figure S5: A selected isolation window ( 11775-11825 m/z) with ECCR applied for BA45 and BA45 with dp20.
